# Identification of Sequence Determinants for the ABHD14 Enzymes

**DOI:** 10.1101/2023.07.30.551196

**Authors:** Kaveri Vaidya, Golding Rodrigues, Sonali Gupta, Archit Devarajan, Mihika Yeolekar, M. S. Madhusudhan, Siddhesh S. Kamat

**Author notes:** To whom the correspondence should be made. These authors contributed equally (Kaveri Vaidya, Golding Rodrigues).

## Abstract

Over the course of evolution, enzymes have developed remarkable functional diversity in catalyzing important chemical reactions across various organisms, and understanding how new enzyme functions might have evolved remains an important question in modern enzymology. To systematically annotate functions, based on protein sequences and available biochemical studies, enzymes with similar catalytic mechanisms and/or aspects of catalysis have been clustered together into an enzyme superfamily. Typically, enzymes within a superfamily have similar overall three-dimensional structures, conserved key catalytic residues, but large variations in substrate recognition sites and residues to accommodate the diverse biochemical reactions that are catalyzed within the superfamily. The serine hydrolases are an excellent example of such an enzyme superfamily, that based on known enzymatic activities and protein sequences, is split almost equally into the serine proteases and metabolic serine hydrolases. Within the metabolic serine hydrolases, are two outlying members, ABHD14A and ABHD14B, that have high sequence similarity, but their functions remained cryptic till recently. While ABHD14A still lacks any functional annotation to date, we recently showed that ABHD14B functions as a lysine deacetylase in mammals. Given their high sequence similarity, automated databases wrongly assign ABHD14A and ABHD14B as the same enzyme, and therefore, annotating functions to them in various organisms maybe problematic. In this paper, we present a bioinformatics study coupled to biochemical experiments, that identifies key sequence determinants for both ABHD14A and ABHD14B, and enables better classification for them. Additionally, we map these enzymes on an evolutionary timescale, and provide a resource in studying these interesting enzymes in different organisms.

## INTRODUCTION

Enzymes are remarkable biological catalysts that greatly facilitate rates of biological reactions*^1, 2^*, ranging from simple hydrolytic reactions to highly complex radical transformations across diverse organisms, and by doing so, often serve as critical metabolic lynchpins. While the repertoire of biochemical reactions catalyzed by enzymes is quite vast and continues to grow steeply, interestingly enough, in comparison, the three-dimensional protein scaffolds on which enzyme active sites are built for catalyzing such reactions remain limited*^3–6^*. Given this fact, it has been hypothesized, that over millions of years of evolution, as the number of catalytic reactions expanded with the complexity in multi-cellular organisms, most enzymes too, correspondingly evolved likely from a common ancestral protein (enzyme) having a defined three-dimensional scaffold, and developed active sites within this protein (enzyme) architecture to catalyze emerging chemical reactions*^7–10^*. Over the past three decades, with the explosion in the publicly available genome sequences and three-dimensional protein structures from different organisms, coupled to high-throughput (experimental) biochemical studies towards annotating function to enzymes, several homologous proteins (enzymes) from diverse organisms that catalyze (overall) different catalytic reactions, but having similar catalytic mechanisms and/or partial chemical reactions and/or aspects of catalysis, have since been grouped together in what is now popularly known as “Enzyme Superfamily”*^4-7, 11^*.

Given the tremendous efforts of numerous research labs, thanks largely the formation of several global consortia, to date, several enzyme superfamilies have been clearly defined, and the biological functions of a significant portion of the distinct members of a majority of the enzyme superfamilies have been elucidated. Amongst such enzyme superfamilies lies the “*Serine Hydrolase*” enzyme superfamily, that is split almost equally in terms of members into two distinct classes: the serine proteases and the metabolic serine hydrolases (mSHs)*^12–14^*. The mSHs sub-group within this superfamily in humans comprises of ∼ 125 members, that catalyze a range of hydrolytic reactions with a consensus ping-pong catalytic mechanism, and deregulation in activities of these enzymes are implicated in numerous human diseases ranging from genetic diseases (e.g. autoimmune conditions and neurological disorders) to metabolic disorders (e.g. dyslipidemia and diabetes) to cancers*^12, 13^*. Of note, most members of the mSH family possess the canonical α/β-hydrolase domain (ABHD)-fold*^15–18^*. Interestingly, while classifying this enzyme family, since several members did not have any known function assigned to it at that time, they were as a monikered ABHDx (x = number) protein.

Based on a comparative sequence phylogenetic analysis*^12, 19^*, at the periphery of the mSH family lie two closely related enzymes, ABHD14A and ABHD14B, that based on their sequence are hypothesized to possess the ABHD-fold, and a conserved catalytic triad (Ser-His-Asp). A major distinction of both these enzymes compared to other mSHs is that the invariant nucleophilic active site serine residue falls within a non-canonical SxSxS motif (canonical motif: GxSxG), hence making them outlying members of this enzyme family. Interestingly, unlike any other mSH, both ABHD14A and ABHD14B were discovered from high throughout screens looking to identify protein interactors of transcription factors. ABHD14A (also known as Dorz1) was found to be a downstream genetic interactor of the transcription factor Zic1, and in mammals, ABHD14A is prospected to have a role in embryonic development of the cerebellum*^20^*. On the other hand, a yeast two-hybrid screen identified ABHD14B (also known as CCG1-interacting protein B, CIB) as a protein interactor of the histone acetyl-transferase (HAT) domain of the general transcription factor TFIID, and was tentatively assigned a role for ABHD14B in transcriptional regulation*^21^*. The same study also reported the three-dimensional structure of ABHD14B, and showed that this enzyme possessed the ABHD-fold*^21^*. It is important to note that both these enzymes were discovered over two decades ago, and no biochemical function has been assigned to either one of them, till recently.

In an effort to functionally annotate these enzymes, we initiated studies on human ABHD14B, and through a series of biochemical and cellular assays, found that this enzyme functions as a novel lysine deacetylase (KDAC)*^22^*. In a follow up study, we found that depleting ABHD14B in mammalian cells and liver of mice results in altered (systemic) glucose metabolism and cellular energetics*^23^*. While the exact biological (protein) substrates and mechanisms of action for this altered metabolic phenotype remain cryptic, our studies have managed to establish a catalytic mechanism for ABHD14B. On the other hand, ABHD14A still remains elusive with regards to any biochemical function. Given their relatively high sequence similarity, we found while studying these enzymes, that across various protein databases, despite being distinct proteins, ABHD14A and ABHD14B were mis-assigned to one another. Therefore, to understand what are minimal sequence determinants for categorizing these enzymes, in this paper, we perform a thorough bioinformatics study on both ABHD14A and ABHD14B, coupled to a few biochemical studies, and present a resource for assigning a given protein sequence as either ABHD14A or ABHD14B. Further, we assess the presence of both these enzymes on the evolutionary time scale, and identify protein sequences in various organisms that correspond to either ABHD14A and ABHD14B, and in doing so, pave the way for better classification of these enzymes in an effort to assigning functions to them in different organisms in the coming years.

## MATERIALS AND METHODS

### Bioinformatics searching and analysis

To determine the prevalence of ABHD14A and ABHD14B protein sequences (and its homologues) across all organisms, PSI-BLAST searches*^24^* were carried out on reference sequences of human ABHD14A (RefSeq: NP_056222.2, Uniprot: Q9BUJ0) and human ABHD14B (RefSeq: NP_001139786.1, Uniprot: Q96IU4), for three iterations, against standard non-redundant databases with an expect threshold of 0.00005 and a maximum return of 5000 hits. To assess the presence or absence of any transmembrane domain(s), all protein sequences from the initial search were submitted to two different transmembrane region prediction software namely CCTOP*^25^* and TMHMM*^26^*. For additional curation of the data, a home-made script was written to verify the presence of the conserved catalytic triad conserved within the searched motifs across the length of the sequence, by carrying out pairwise global alignments using the Needleman-Wunsh algorithm*^27^*. For the final resulting of the datasets, a single protein sequence was taken from each representative organism, and subjected to a multiple sequence global alignment using MAFFT*^28^*, the alignments were further used in MEGA-X*^29^* to build maximum likelihood trees, and the trees were visualized and annotated in iTol*^30^*. The fully conserved residues for both ABHD14A and ABHD14B were identified using pymsaviz (pypi.org/project/pymsaviz/) and Bio.Align (biopython.org/docs/1.76/api/Bio.Align.html) packages in Python, and the functionally conserved residues for both enzymes were identified by manual inspection of sequences.

### Cloning, Expression and Purification of ABHD14B Mutants

The human *abhd14b* gene*^22^* was synthesized as a codon-optimized construct (from Genscript) for expression in *E. coli* and, further sub-cloned into the pET-45b(+) vector (Millipore) with a N-terminal (His)_6_-tag. For generating single-point mutants, site directed mutagenesis was performed using Pfu DNA polymerase as per manufacturer’s instructions (Promega, Cat. # M7741). The resulting sequence verified plasmids having the desired mutation were individually transformed into *E. coli* BL21(DE3) competent cells, following which, a single colony was inoculated into 5 mL of Luria Bertani (LB) medium containing ampicillin (final concentration in media was 100µg/mL) and grown overnight (12-14 hours) in an incubator with a temperature at 37°C with constant shaking. This primary inoculum was transferred to 1L of the same medium, and grown in an incubator with a temperature at 37°C with constant shaking until the absorbance at 600 nm reached ∼ 0.6. At this time-point, the temperature of the incubator was reduced to 18°C, and the protein expression was induced by adding 500 μM of isopropyl-β-D-1-thiogalactopyranisode to the culture. Upon induction of protein expression, the bacterial culture was grown for an additional 16 – 18 hours at 18°C, following which, the cells were pelleted by centrifugation at 6000g for 20 minutes and stored at −80°C until further use. The ABHD14B overexpression was assessed by SDS-PAGE analysis and only upon confirming robust protein overexpression, the resulting cell pellets were used for further processing.

The ABHD14B mutants were purified by Ni-NTA affinity chromatography using a His-Trap High Performance column (GE Heathcare, Cat #17-5248-02) as per manufacturer’s instructions. Briefly, a cell pellet (with confirmed overexpression of the ABHD14B mutant) was thawed and re-suspended in 45 mL of Binding Buffer (50 mM Tris, 10 mM imidazole at pH 8) at 4°C. Cells were lysed using a Vibra Cell VCX 130 probe sonicator (Sonics) at 60% amplitude for 30 minutes with 2 seconds “ON” and 3 seconds “OFF” cycle on ice. The resulting homogenate was centrifuged for 30 minutes at 30,000g at 4°C to separate the soluble and insoluble fraction, and the soluble fraction (that contains ABHD14B mutants) was applied to a pre-equilibrated His-Trap High Performance column. The desired ABHD14B mutant was eluted using an increasing imidazole gradient (50 mM to 500 mM) to the binding buffer, and the purity of the various collected fractions of the eluted ABHD14B mutant was confirmed by SDS-PAGE analysis. The fractions containing the pure ABHD14B mutant were pooled, concentrated in 50 mM Tris (pH 8) using a 10 kDa molecular weight cut off filter (Millipore), and ∼ 4 mL of this concentrated ABHD14B mutant was loaded onto a pre-equilibrated HiLoad^TM^16/60 Superdex200 (GE Healthcare) gel filtration column. The fractions collected from this size exclusion chromatography column were assessed for the purity of the desired ABHD14B mutant by SDS-PAGE analysis, and those containing the desired pure ABHD14B mutant were pooled, concentrated as described earlier, aliquoted (10 μL aliquots), flash-frozen using liquid nitrogen, and subsequently stored at −80°C till further use.

### Gel Based <u>A</u>ctivity <u>B</u>ased <u>P</u>rotein <u>P</u>rofiling (ABPP) Experiments

All gel-based ABPP assays were performed as reported earlier*^22, 31, 32^*. In these assays, the total reaction volume was 50 μL, while the final protein and activity probe (fluorophosphonate-rhodamine, FP-rhodamine) concentrations were 5 μM each. All samples were resolved on a 12.5% SDS-PAGE gel, and the activities of the mutants were visualized using a iBright1500 (Invitrogen) gel documentation system.

### Colorimetric Substrate Hydrolysis Assays

All colorimetric substrate hydrolysis assays with *p-*nitrophenyl-acetate (pNP-acetate) were performed as reported earlier*^22^* in 50 mM Tris at pH 8. All substrate assays were performed in a final reaction volume of 250 μL, with a final concentration of 10 µM and 100 µM for the ABHD14B mutant and pNP-acetate respectively, in biological triplicate to ensure reproducibility. For assessing the relative activities of the ABHD14B mutants, 500 second time-point was chosen, as this colorimetric assay was found to be linear with a good signal to noise ratio at this time point.

### Data Plotting

All data presented in this paper was plotted using the GraphPad Prism 9 (version 9.5.1) software for MacOS-X. Unless otherwise mentioned, all data in presented as the mean ± the standard deviation for at least three independent experiments (biological replicates).

## RESULTS AND DISCUSSION

### Identification of ABHD14B Sequences

The three-dimensional structure of human ABHD14B has shown that this metabolic serine enzyme has the ABHD-fold, and possess a conserved catalytic triad, consisting of Ser-111, Asp-162 and His-188 (**Figure 1**)*^21^*. In addition to this, a bioinformatics survey of various ABHD-proteins from the metabolic serine hydrolase family have identified two conserved sequence motifs for (mammalian) ABHD14Bs, namely the nucleophilic motif (consisting of a SxSxS motif within the VVI<u>SPSLS</u>GMY sequence) and an acyltransferase motif (consisting of the HxxxxD motif within the GAG<u>HPCYLD</u>KPE sequence) towards the C-terminal end of the protein (**Figure 1**)*^15^*. From the 5000 hits obtained from using human ABHD14B (RefSeq: NP_001139786.1, Uniprot: Q96IU4) as a query sequence for search, the ABHD14B nucleophilic motif was identified in 1012 sequences (allowing for up to two mismatches), of which 792 sequences were perfect matches. Amongst the 5000 hits, the ABHD14B acyltransferase motif was identified in 1265 sequences (allowing for up to two mismatches), with 217 sequences being perfect matches. Of note, the catalytic triad was identified in 2929 sequences. Based on the above information, in our study, a protein sequence was classified as ABHD14B, if it possessed the catalytic triad, the ABHD14B nucleophile motif (allowing for up to two mismatches), and the ABHD14B acyltransferase motif (allowing for up to two mismatches). Based on this filtering criteria, overall, we identified 847 ABHD14B sequences identified from 697 organisms, of which 197 ABHD14B sequences identified from 130 organisms were perfect matches to the aforementioned filtering criteria (**Supplementary Table 1**).

**Figure 1.**
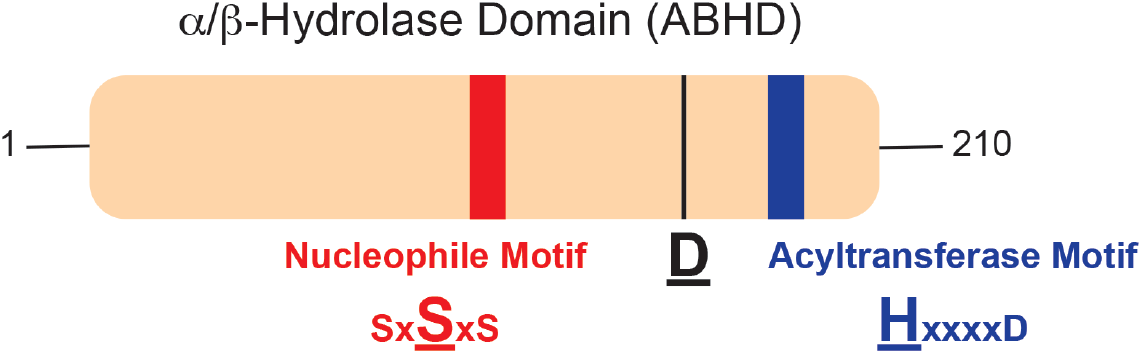
Schematic representation of the human ABHD14B structure. Based on available literature, ABHD14B is predicted to contain the catalytic triad consisting of Ser-Asp-His, a nucleophilic motif that contains the active site serine residue as part of a SxSxS sequence, and an acyltransferase motif that contains the HxxxxD sequence, all within an overall ABHD-fold.

### Phylogenetic Classification of ABHD14B Sequences

From the filtering of protein sequence obtained from the database searches, we identified 847 ABHD14B sequences, and found that these protein sequences came from 697 organisms. Upon manual curation of this data, it was clear, that quite a few organisms (especially from class *Mammalia* and *Aves*) possessed more than one isoform of ABHD14B. Hence, the total number of ABHD14B sequences was significantly greater than organisms identified from our search (**Supplementary Table 1**). Nonetheless, we chose the longest ABHD14B sequence from any particular organism to perform a phylogenetic (evolutionary) analysis for ABHD14B, and found that ABHD14B was almost exclusively confined to phylum *Chordata* (∼ 99.7 %, 695 organisms of the 697 organisms identified) (**Figure 2A**). Amongst the phylum *Chordates*, the ABHD14B sequences were found most in class *Aves* (Birds) (∼ 45%, 313 organisms of the 697 organisms identified), *Mammalia* (Mammals) (∼ 27%, 189 organisms of the 697 organisms identified) and *Actinopterygii* (Bony Fish) (∼ 22%, 156 organisms of the 697 organisms identified), with a smaller representation seen in *Reptilia* (Reptiles) (∼ 5%, 33 organisms of the 697 organisms identified) (**Figure 2B**). Based on the phylogenetic analysis, the avian ABHD14B protein sequences were most closely related to the mammalian ABHD14B sequences, while those class *Actinopterygii* and *Reptilia* were most closely related to one another, and together had a closer sequence relation to mammalian ABHD14Bs than the avian ABHD14Bs (**Figure 2A**). It was interesting to note that of the 697 organisms possessing ABHD14B sequence, 263 organisms possessed ABHD14A sequence (∼ 38% of the total organisms) (**Figure 2**) (discussed in “*Phylogenetic Classification of ABHD14A Sequences*” section).

**Figure 2.**
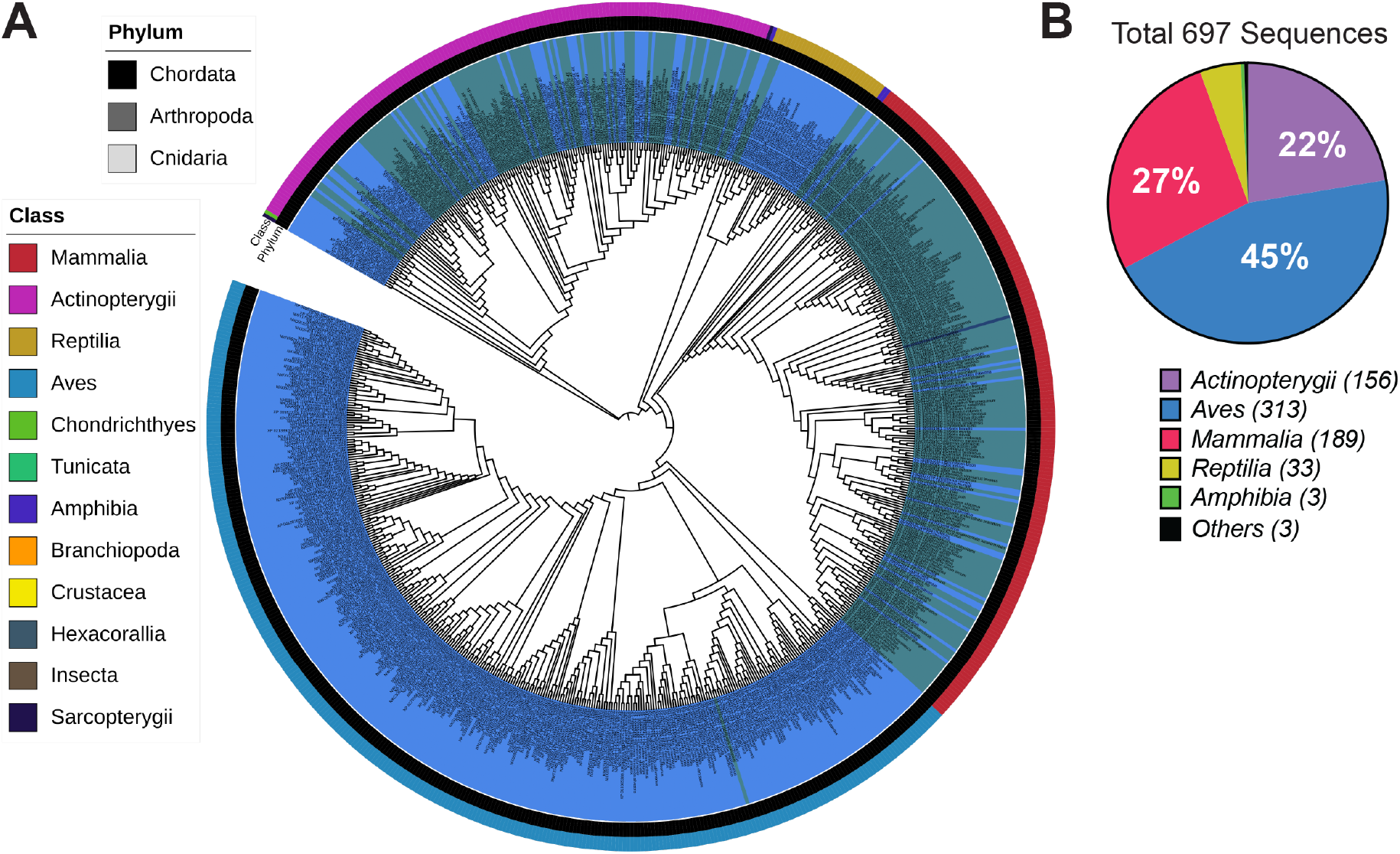
Phylogenetic analysis of ABHD14B sequences. (**A**) Phylogenetic tree representing the identified ABHD14B sequences from 697 organisms. The outermost and middle circular coloring denotes the Class and Phylum to which a sequence belongs (see associated legends within the figure), while the innermost circular coloring (blue or green) denotes whether an organism contains a sequence only for ABHD14B (blue) or for both ABHD14B and ABHD14A (green). (**B**) Pie-chart analysis representing the data from the phylogenetic tree for the various classes of organisms, mainly from *Chordates* that contain a ABHD14B sequence. This analysis shows that class *Aves*, *Mammalia* and *Actinopterygii* contain the most sequences for ABHD14B within the *Chordates*.

### Conservation of Residues within the ABHD14B Sequences

To assess the overall conservation in protein sequence across all the 697 ABHD14B sequences identified from different organisms following database searches and subsequent phylogenetic analysis, we performed a multiple sequence alignment analysis on them (**Supplementary File 1**). From this analysis, we found that across all the 697 ABHD14B sequences, 86 residues were highly conserved (present at a frequency of > 90% in all the 697 ABHD14B sequences at the defined position), suggesting that across all organisms, the overall sequence conservation (and thereby identity) is fairly high (∼ 40%). Additionally, upon closer inspection, we found that within a particular class (e.g. *Mammalia*), the extent of sequence conservation is significantly higher (∼ 80%) (**Supplementary File 1, Supplementary Table 1**). Besides the overall conserved residues, we found that an additional 20 residues were functionally conserved (presence of similar amino-acid type at that position; e.g. Glu and Asp are functionally conserved residues) across all the 697 ABHD14B sequences, suggesting that the realistic sequence conservation across all organisms for ABHD14B was ∼ 50% (**Supplementary File 1, Supplementary Table 1**).

Next, we mapped all the conserved residues (both absolute conserved and functionally conserved) on the available three-dimensional structure of ABHD14B (PDB: 1imj), and found not surprisingly that these conserved residues were clustered around the enzyme active site and the cleft adjoining this active site where the putative substrates are predicted to bind (**Figure 3**). Further, we also found from this structural analysis, that the residues present on the flexible loops involved in substrate recognition were also highly conserved. Next, previous modeling studies from our lab towards understanding the mechanism of this enzyme*^22^*, revealed that ABHD14B has two tunnels that seem to converge on the active site serine residue: (i) The putative peptide (substrate # 1) tunnel; and (ii) The coenzyme A (substrate # 2) tunnel (**Figure 4**). It is interesting to note that the entrance to both these tunnels have highly conserved positively charged residues presumably gating the entrance of substrates to the active site (for the human ABHD14B, Arg42 and Lys141 at the entrance of the coenzyme A and peptide tunnel respectively). Further, we noticed that the residues that comprise of these tunnels also show high degree of conservation, and of note, we found that the coenzyme A tunnel comprises of several hydrophobic residues, presumably to exclude water from the active site, to prevent the hydrolysis of the acetyl-enzyme intermediate*^22, 23^*.

**Figure 3.**
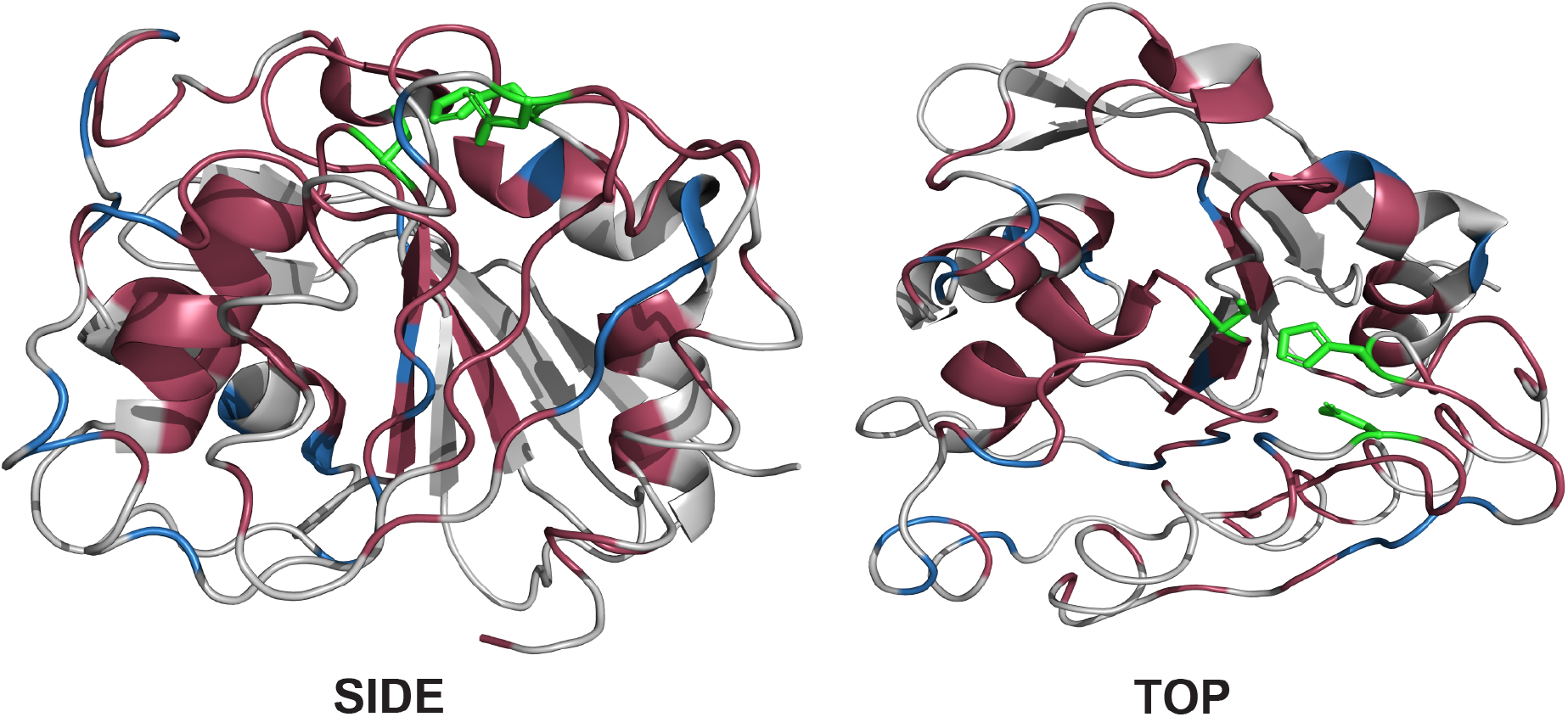
Mapping conserved residues on the ABHD14B structure. The residues colored in red are those that are absolutely conserved, while those shown in blue are functionally conserved based on the multiple sequence alignment analysis across all the 697 sequences of ABHD14B from different organisms. The catalytic triad residues that mark the active site are shown in green.

**Figure 4.**
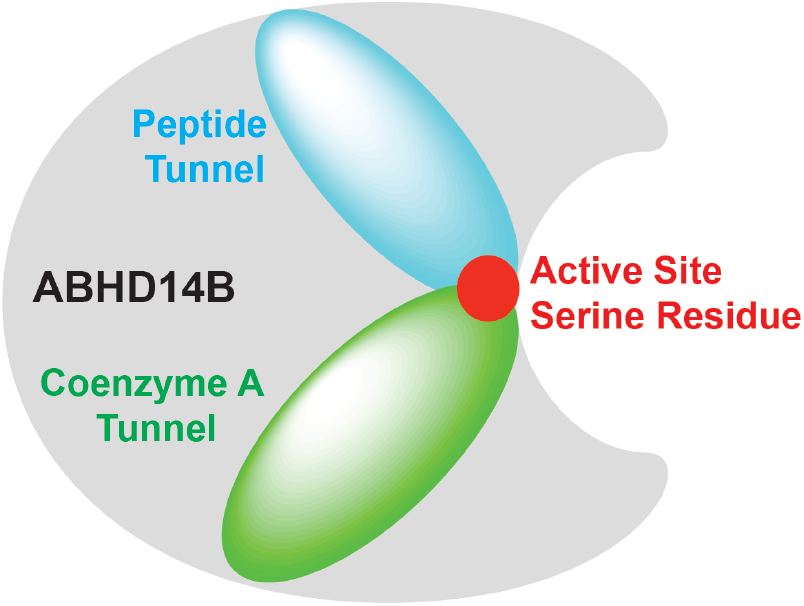
Cartoon representation of the substrate tunnels within ABHD14B. The three-dimensional structure of human ABHD14B suggests that this enzyme has two tunnels for the two substrates that converge on the active site serine residue.

### Biochemical Characterization of ABHD14B mutants

The bioinformatics analysis helped us identify conserved residues across the different ABHD14B sequences from various organisms. Next, using biochemical assays, we wanted to validate some of these bioinformatics findings, and assess the effect of mutating some key conserved residues on ABHD14B activity. Towards this, we chose to assay the recombinant human ABHD14B, as we and others have previously studied this cryptic enzyme and a three-dimensional structure is also available for it*^21–23^*. For assaying the various human ABHD14B mutants, we chose two previously established biochemical assays, namely the gel-based ABPP chemoproteomic platform and pNP-acetate based colorimetric substrate hydrolysis assay*^22^*. Using an activity based probe, FP-rhodamine in this case, the gel-based ABPP assay reports on the nucleophilicity of the active site serine residue, and by doing so, serves as a proxy for the activity of an enzyme*^22, 33^*. On the other hand, the pNP-acetate based colorimetric substrate hydrolysis assay serves as a direct readout for the overall catalytic ability of the enzyme to turn over an acetylated substrate surrogate*^22^*.

First, we made alanine mutants for the residues from the catalytic triad, i.e. S111A, D162A and H188A, and found from gel-based ABPP assays that these mutants were unable to react with the activity probe (FR-rhodamine), and show signals on the activity gel (**Figure 5A**). In addition to this, relative to wild type (WT) ABHD14B, all the three mutants also showed > 50% loss in the ability to turn over pNP-acetate, with S111A having the most diminished activity as expected (**Figure 5B**). Second, we focused on the two gate-keeping residues of the tunnels, namely R42 (coenzyme A tunnel) and K141 (peptide tunnel), and made alanine mutants for both these residues. Upon assaying them, we found that relative to WT ABHD14B, there was no effect in both the assays for the R42A mutant (**Figure 5A, 5B**). Interestingly however, while relative to WT ABHD14B, the K141A mutant did not show any appreciable change in the gel-based ABPP assay (**Figure 5A**), mutation to K141 did result in > 50% reduction in its ability to turn over pNP-acetate (**Figure 5B**). This result strongly suggests that K141 plays an important role in the orientation of the (peptide) substrate for optimal catalysis. A similar role for R42 also cannot be excluded just based on these assays, since we are not testing the enzyme’s ability to bind coenzyme A in either biochemical assay, and a further biochemical analysis on this mutant with coenzyme A may be needed to ascertain this role. Third, we wanted to assess the role that D193 plays during this catalysis, as this residue is part of the invariant HxxxxD acyltransferase motif found in all ABHD14B sequences. Interestingly, when we assayed the D193A mutant, we found that while it had almost WT ABHD14B level of pNP-acetate hydrolysis activity (**Figure 5B**), the nucleophilicity of the active site serine was significantly diminished, as evidenced from its inability to react with the activity probe (FP-rhodamine) from the gel-based ABPP assays (**Figure 5A**). Based on known mechanisms for acyltransferases, by regulating the nucleophilicity of S111, presumably due to its interactions with H188 as part of the HxxxxD motif, D193 might be have an important role to play in the transfer of the acetyl-group from the acetyl-enzyme intermediate to coenzyme A (second-half of enzyme cycle)*^22, 23^*. This premise also needs to be further tested with more intricate biochemical assays using coenzyme A and the D193A mutant.

**Figure 5.**
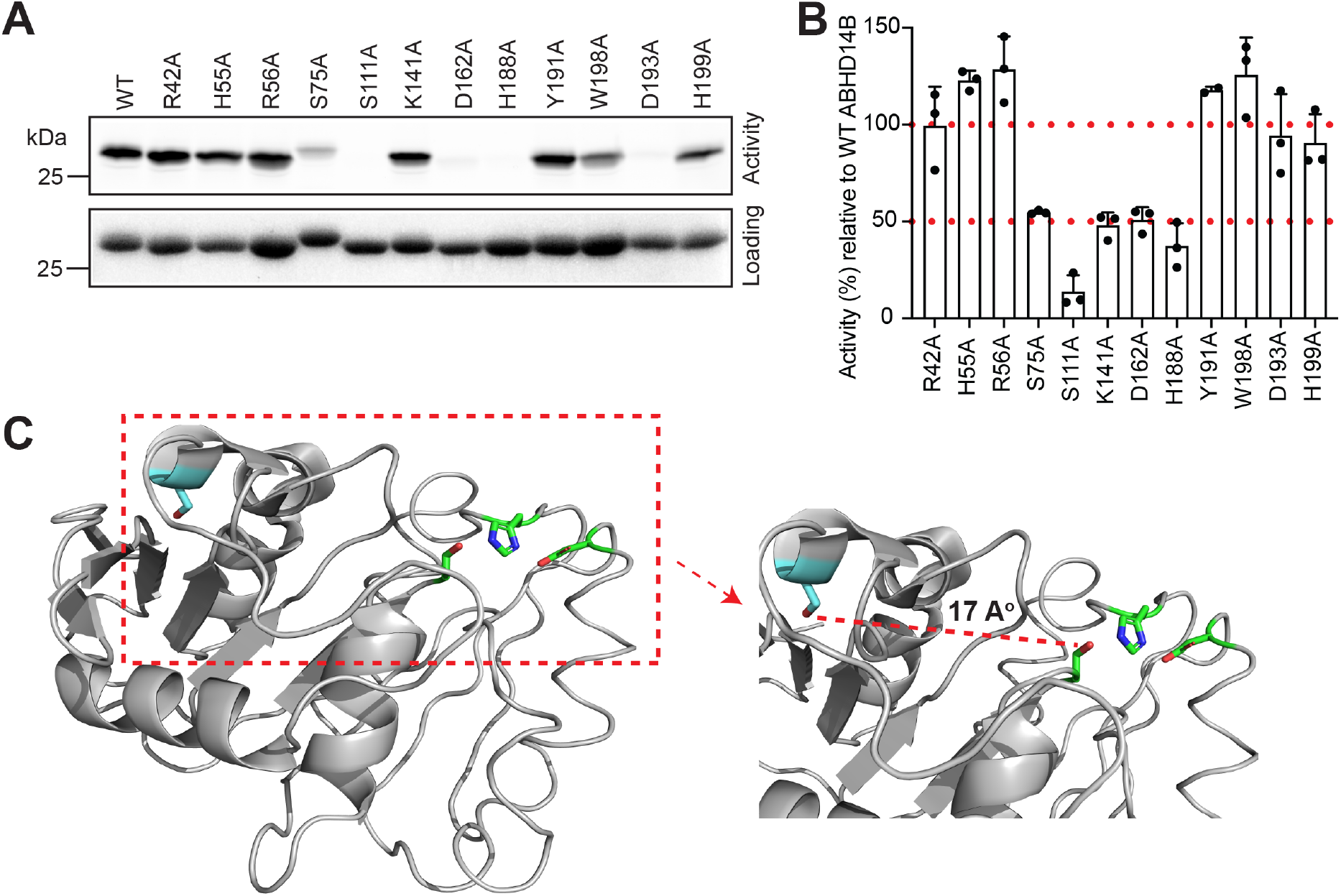
Biochemical characterization of ABHD14B mutants. (**A**) Representative gel showing the activity of the various mutants generated for human ABHD14B in a gel-based ABPP assay. In this assay, the protein and probe concentrations were 5 μM each. The top-gel shows in-gel fluorescence, while the bottom gel shows equal protein loading by Coomassie staining. This experiment was repeated three independent times (biological replicates) with reproducible results each time. (**B**) pNP-acetate substrate hydrolysis assay showing the activity of various mutants in this colorimetric assay. The bar data denotes mean ± standard deviation for the individual data points (three biological replicates), relative to WT ABHD14B (control). In this assay, the protein and pNP-acetate concentrations were 10 and 100 μM respectively. (**C**) The three-dimensional structure of human ABHD14B showing the position of S75 (shown in blue) relative to the catalytic triad (shown in green). The zoomed image of this, shows that S75 is approximately 17 A° away from the active site S111 residue on a flexible loop.

Next, we wanted to check if mutations to residues within a protein tunnel had any effect on ABHD14B activity, and we chose residues from the coenzyme A tunnel, as previous structural modeling studies from our lab had speculated some roles for different residues within this tunnel*^22^*. Towards this, we made alanine mutants for H55, R56, Y191, W198 and H199, that are conserved, and predicted to the neutralize the negative charges on phosphates (H55, R56) or π-stack with and stabilize the nucleotide region of coenzyme A (Y191, W198, H199). However, we found that relative to WT ABHD14B, the single point alanine mutants of the aforementioned residues (H55A, R56A, Y191A, W198A, and H199A) did not have much effect on ABHD14B activity in either assay (**Figure 5A, 5B**). We also found that these mutants did not have any defects in their ability to bind coenzyme A (data not shown). This result suggests that given the tight regulation in this enzymatic activity and the rigid overall structure of this tunnel, perhaps multiple residues might need to be mutated simultaneously to see any effect on ABHD14B activity. Lastly, we found from the bioinformatics and multiple sequence alignment analysis that S75 (in humans) was a universally conserved residue in ABHD14B sequences from all organisms (**Supplementary File 1**). This was an intriguing finding, as based on the three-dimensional structure of human ABHD14B, S75 resides on a flexible loop that is present ∼ 17 A° away from the active site, and it does not seem to have any particular predicted role in the enzyme catalytic cycle (**Figure 5C**). However, we found that upon mutation of this residue, the S75A mutant had significantly diminished activity in both the gel-based ABPP (**Figure 5A**) and pNP-acetate hydrolysis (**Figure 5B**) assay. Several post-translational modification (PTM) prediction databases/algorithms seem to suggest that ABHD14B might be phosphorylated, and that perhaps this protein phosphorylation might have physiological implications in different biological contexts. Though only speculative yet, given the activity profile seen from the assays done with the S75A mutant, it might be possible that through its PTM, S75 might be allosterically regulating ABHD14B activity. Overall, the information from the conserved residues has helped confirm known findings and open new avenues for residues that may be tested to probe the enzymatic mechanism of ABHD14B further.

### Identification of ABHD14A Sequences

Owing to its high sequence similarity to ABHD14B, for a given organism, various bioinformatics studies have predicted that the three-dimensional structure of ABHD14A also has the ABHD-fold, and possess a conserved catalytic triad, consisting of Ser-171, Asp-222 and His-249 in the human ABHD14A sequence (**Figure 6**). Like ABHD14B, bioinformatics studies with other ABHD-proteins from the metabolic serine hydrolase family have identified a conserved nucleophilic motif (consisting of a SxSxS motif within the VLV<u>SPSLS</u>GHY sequence) (**Figure 6**)*^15^*. Of note, bioinformatics studies suggest that ABHD14A lacks the acyltransferase motif seen in ABHD14B, and unlike ABHD14B, surprisingly has an integral membrane domain consisting of ∼ 30 – 40 amino acids that form an anchoring α-helical sequence at the N-terminal end of the protein (**Figure 6**). Of the 5000 hits obtained from using human ABHD14A (RefSeq: NP_056222.2, Uniprot: Q9BUJ0) as a query sequence for search, the nucleophilic motif for ABHD14A was identified in 733 sequences (allowing for up to two mismatches), of which 134 sequences were perfect matches, while the catalytic triad was identified in 2624 sequences. From the 5000 hits obtained from the initial search, the two transmembrane prediction software, CCTOP and TMHMM predicted transmembrane regions in 1069 and 1084 sequences respectively, of which a total of 1010 sequences were common to both prediction software, while 129 were identified by a single software, leading to a complete set of 1139 protein sequences with an identified transmembrane region. Based on the above information, in our study, a protein sequence was classified as ABHD14A, if it possessed the catalytic triad, the ABHD14A nucleophile motif (allowing for up to two mismatches), and had a transmembrane domain. Based on this filtering criteria, overall, we identified 483 ABHD14A sequences identified from 312 organisms, of which 116 ABHD14A sequences identified from 52 organisms were perfect matches to the aforementioned filtering criteria (**Supplementary Table 2**).

**Figure 6.**
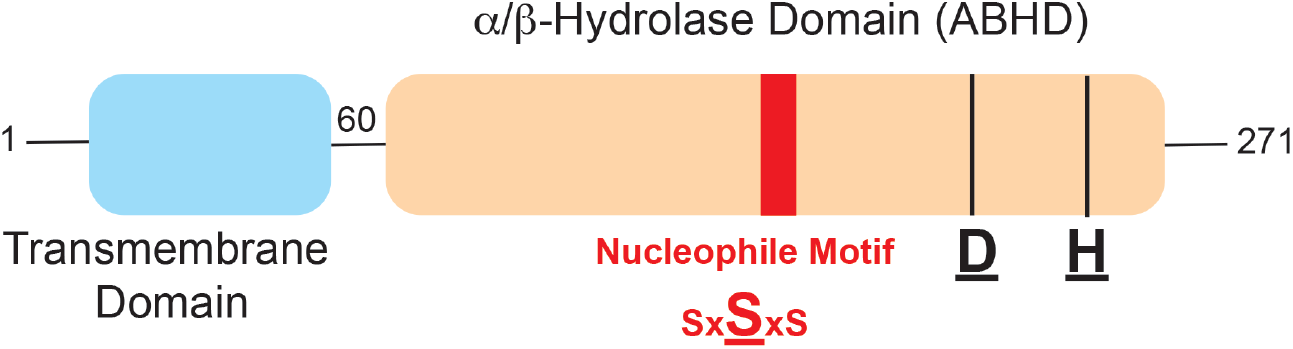
Schematic representation of the human ABHD14A structure. Based on available literature, ABHD14A is predicted to contain the catalytic triad consisting of Ser-Asp-His, and a nucleophilic motif that contains the active site serine residue as part of a SxSxS sequence within an overall ABHD-fold. Additionally, ABHD14A contains a N-terminal domain that is predicted to anchor it to the cellular membranes.

### Phylogenetic Classification of ABHD14A Sequences

Like ABHD14B, upon applying the aforementioned filtering criteria for the shortlisted ABHD14A sequences obtained from various databases, we identified 483 ABHD14A sequences that came from 312 organisms. Like ABHD14B, upon manual curation of this data, it was clear, that several organisms (especially from class *Mammalia* and *Actinopterygii*) possessed more than 1 isoform of ABHD14A, and therefore, there was a disconnect between the total ABHD14A sequences and the organisms identified from our search (**Supplementary Table 2**). Like the earlier ABHD14B analysis, we chose the longest ABHD14A sequence from any particular organism to perform a phylogenetic (evolutionary) analysis for ABHD14A, and found that ABHD14A was largely confined to phylum *Chordata* (∼ 97.5%, 304 organisms of the 312 organisms identified), with some representation from the *Arthropoda* and *Cnidaria* phyla (**Figure 7A**). Amongst the phylum *Chordates*, the ABHD14A sequences were predominantly found in class *Mammalia* (Mammals) (∼ 58%, 182 organisms of the 312 organisms identified) and *Actinopterygii* (Bony Fish) (∼ 37%, 114 organisms of the 312 organisms identified), with a much smaller representation seen in *Reptilia* (Reptiles) (∼ 2%, 6 organisms of the 312 organisms identified) (**Figure 7B**). The phylogenetic analysis showed that within the *Chordates*, the ABHD14A protein sequences from class *Actinopterygii* and *Mammalia* were most closely related to one another, and together were related to ABHD14A sequences from the class *Reptilia* (**Figure 6**). Of the 312 organisms that we identified possessing a ABHD14A sequence, 263 organisms also possessed a ABHD14B sequence, an overall fraction (∼ 84% of the total organisms) far greater than ∼ 38% observed from the ABHD14B analysis (**Figure 7C, Supplementary Tables 1, 2**).

**Figure 7.**
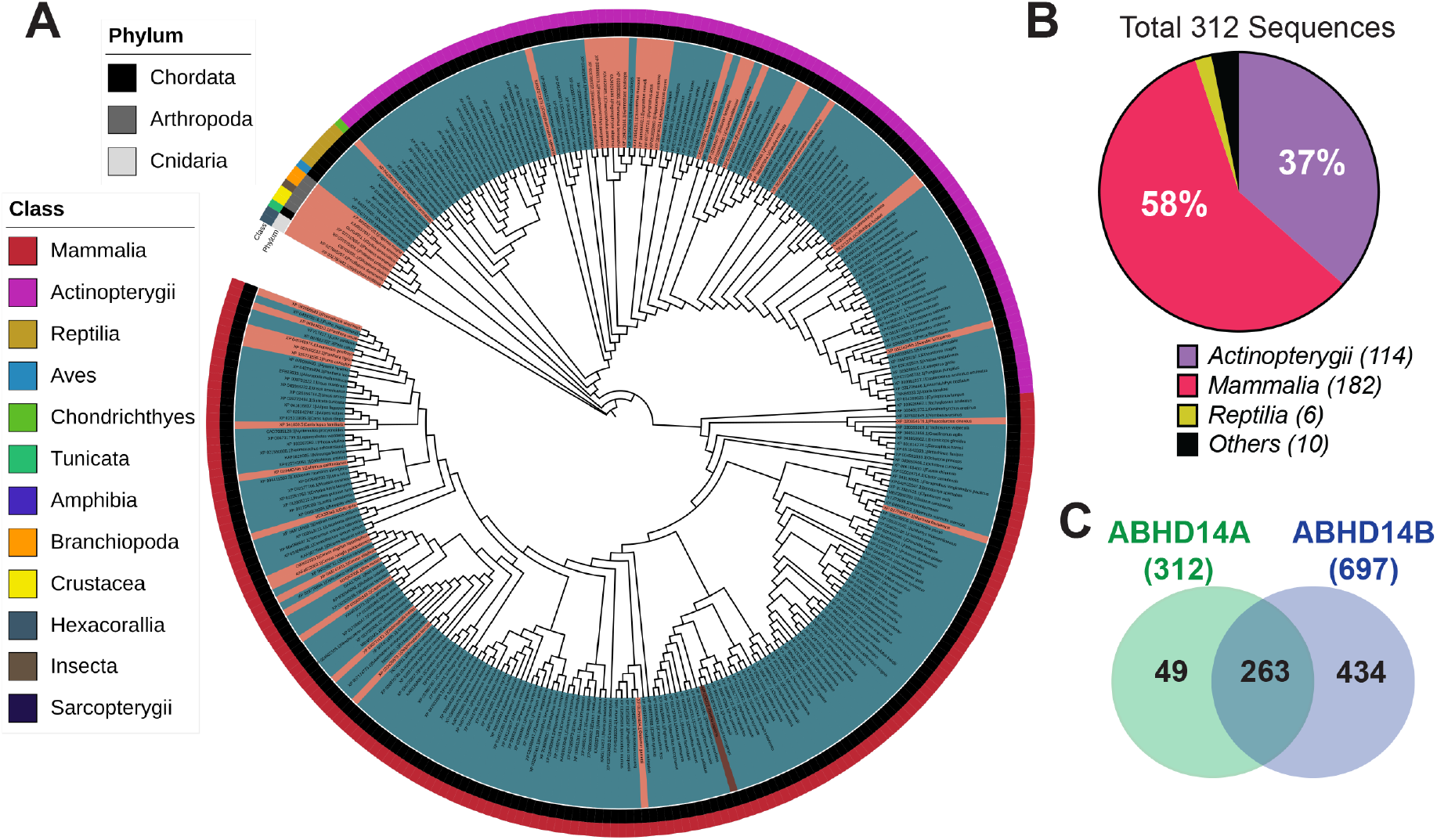
Phylogenetic analysis of ABHD14A sequences. (**A**) Phylogenetic tree representing the identified ABHD14A sequences from 312 organisms. The outermost and middle circular coloring denotes the Class and Phylum to which a ABHD14A sequence belongs (see associated legends within the figure), while the innermost circular coloring (orange or teal) denotes whether an organism contains a sequence only for ABHD14A (orange) or for both ABHD14B and ABHD14A (teal). (**B**) Pie-chart analysis representing the data from the phylogenetic tree for the various classes of organisms, mainly from *Chordates* that contain a ABHD14A sequence. This analysis shows that class *Mammalia* and *Actinopterygii* contain the most sequences for ABHD14A within the *Chordates*. (**C**) Venn diagram showing the overlapping number of organisms that contain ABHD14A and ABHD14B sequences amongst those identified from the bioinformatics analysis.

The most striking and surprising aspect of this phylogenetic analysis for ABHD14A was that based our filtering criteria, we failed to identify any ABHD14A sequences from class *Aves* within the *Chordates* (**Figure 7A, 7B**). This runs counter to the fact that *Aves* possessed the highest number of sequences for ABHD14B, suggesting that there might be three broad possibilities: (i) Perhaps our filtering criteria for identification of ABHD14A were too stringent; and/or (ii) *Aves* did not possess sequences for ABHD14A (lost during evolution); and/or (iii) ABHD14A sequences possessed by *Aves* were very distinct from ABHD14A sequences identified from other organisms. To exclude, possibility (i), we re-searched the 5000 hits, lowering mismatch threshold to three and four mismatches, and were still unable to identify any ABHD14A sequences from *Aves*. Our current studies therefore, cannot exclude the possibilities (ii) and (iii), and further bioinformatics analysis for ABHD14A sequences specific to class *Aves* will be needed to resolve this preliminary yet interesting finding.

### Conservation of Residues within the ABHD14A Sequences

Like the previously described analysis done for ABHD14B, to assess the overall conservation in protein sequence for ABHD14A, we performed a multiple sequence alignment analysis on all the 312 sequences identified for ABHD14A from different organisms (**Supplementary File 2**). Here, we found that from the 312 ABHD14A sequences, 99 residues were highly conserved (present at a frequency of > 90% in all the 312 ABHD14A sequences at the defined position), suggesting that across all organisms, like ABHD14B, the overall sequence conservation (and thereby identity) is fairly high (∼ 40%). Additionally, like ABHD14B, upon closer inspection, for ABHD14A sequences, we found that within a particular class (e.g. *Mammalia*), the extent of sequence conservation was significantly higher (∼ 80%) (**Supplementary File 2, Supplementary Table 2**). Besides the conserved residues, we found that 50 residues were functionally conserved for the 312 ABHD14B sequences, suggesting that the overall sequence conservation across all organisms for ABHD14B was realistically ∼ 60% (**Supplementary File 2, Supplementary Table 2**).

Next, like we did for ABHD14B, we wanted to map the conserved residues identified for ABHD14A to a three-dimensional protein structure. However, in the absence of any available ABHD14A structure, we used a predicted structure of human ABHD14A (Uniprot ID: Q9BUJ0) that was generated by the AI algorithm/system of the AlphaFold Protein Structure Database*^34, 35^*. To ensure that this structure was useful for further analysis, we first overlaid this AlphaFold generated structure of human ABHD14A, with the crystal structure of human ABHD14B (PDB: 1imj). From this structural analysis, we found an almost perfect overlay of the ABHD-fold portion of ABHD14A with the overall structure of ABHD14B (**Figure 8**). In fact, the catalytic triad that comprises of the active site of both these enzymes, showed near identical orientations (**Figure 8**), and gave us confidence that the overall predicted structure of human ABHD14A would be useful for our structural analysis. Interestingly, the N-terminal region of the predicted human ABHD14A structure showed both an extended disordered stretch and an α-helical region that presumably anchors this enzyme to a cellular membrane (**Figure 8**). Finally, we mapped all the conserved residues on the AlphaFold predicted structure of human ABHD14A, and like the ABHD14B analysis, found that these conserved residues (both absolute conserved and functionally conserved) were clustered around the enzyme active site (catalytic triad) and the cleft adjoining this active site where the putative substrates of this enzyme are predicted to bind (**Figure 9**). Further, we also found from this structural analysis, that the residues present on the α-helical region at the N-terminus of the protein that are predicted to be involved in membrane anchoring of ABHD14A were also highly conserved (**Figure 9**). This is interesting, because unlike the cellular localization of ABHD14B (cytosolic and nuclear)*^22^*, ABHD14A might be localized to the plasma membrane or microsomal components (e.g. Golgi, endoplasmic reticulum, peroxisome), and may have functions distinct from ABHD14B. While only speculative based on the bioinformatics data, this premise however needs to be experimentally validated. Also of note and biological importance, is that despite the high overall sequence similarity (∼ 40 – 50%) of ABHD14A and ABHD14B within a particular organism that possess both enzymes (e.g. humans), it is interesting to note that ABHD14A lacks the acyltransferase motif (HxxxxD). This implies that unlike the ABHD14B, perhaps ABHD14A performs a function distinct from the known acyltransferase-type activity of ABHD14B, and in the coming years, finding a biological activity/function to ABHD14A might be interesting from an evolutionary and phylogenetic standpoint.

**Figure 8.**
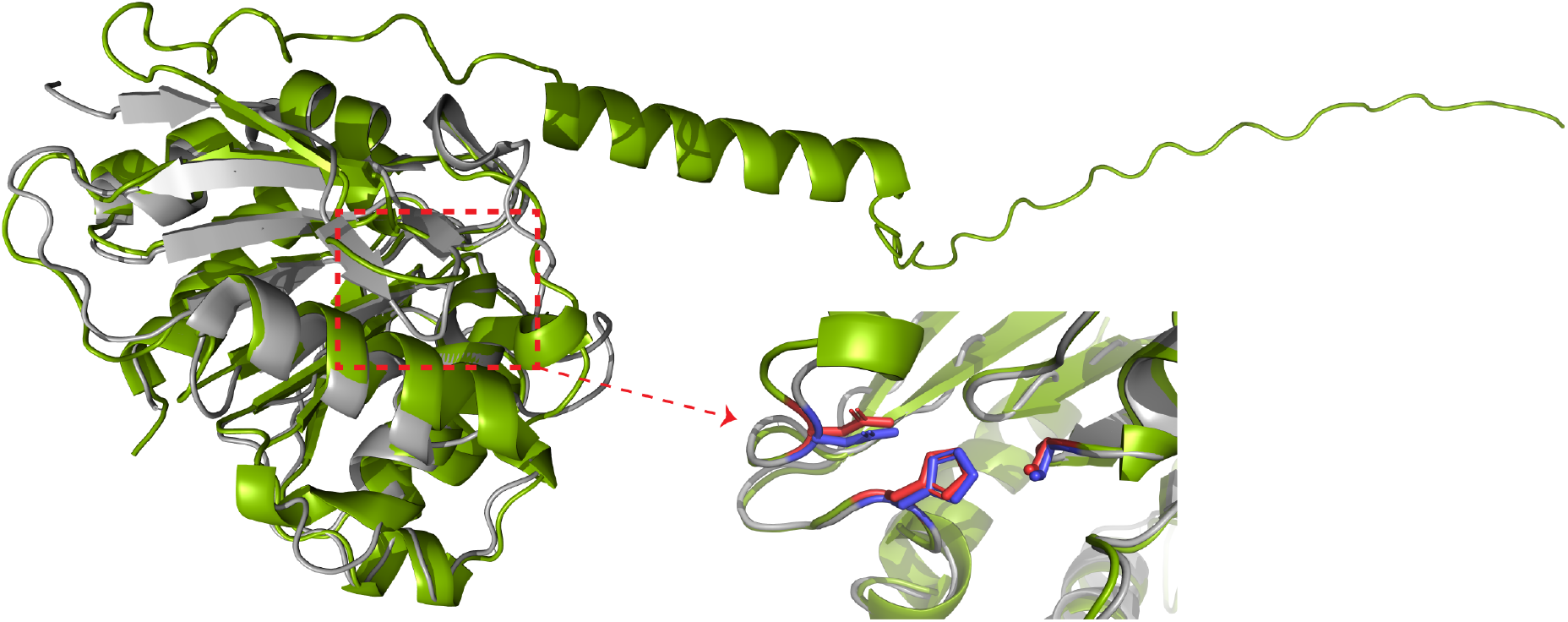
Overlay of the human ABHD14A and human ABHD14B structures. Superimposition of the AlphaFold predicted structure of human ABHD14A (Uniprot ID: Q9BUJ0) (green structure) with the three-dimensional crystal structure of ABHD14B (PDB: 1 imj) (gray structure) showing an almost perfect overlay of the ABHD-fold region of both proteins. A zoomed image of the catalytic triad also shows that these residues are perfectly aligned in both structures (blue residues for ABHD14B and red residues for ABHD14A). The ABHD14A predicted structure shows an additional N-terminal region that comprises of an α-helix, that is responsible for membrane anchoring of this enzyme.

**Figure 9.**
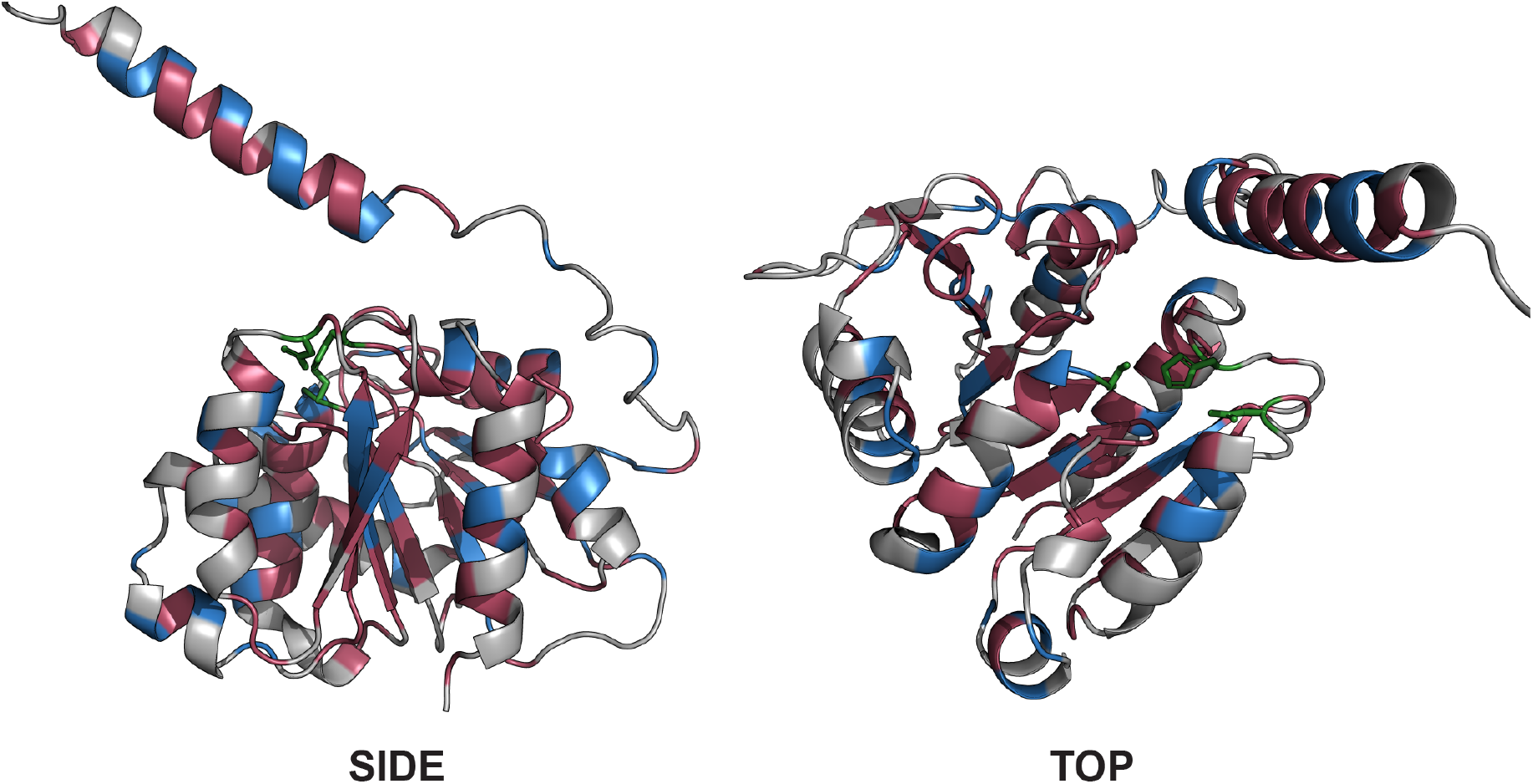
Mapping conserved residues on the ABHD14A structure. The residues colored in red are those that are absolutely conserved, while those shown in blue are functionally conserved based on the multiple sequence alignment analysis across all the 312 sequences of ABHD14A from different organisms. The catalytic triad residues that mark the active site are shown in green.

## CONCLUSION

The advent of genome sequencing technologies has resulted in a deluge of available protein sequences, and mis-annotations of protein functions across public databases are increasingly propagating due to the over-reliance of automated tools of protein function assignment*^11^*. To overcome this, several efforts have been undertaken to tie in high-throughput bioinformatics analysis to experimental outputs to ensure proper validation of functions*^36^*. Realizing that this is an important problem especially in an era of high volumes of data and AI tools for interpreting them, here, we focused on development of a framework to identify and classify two outlying enzymes from the metabolic serine hydrolase family*^12^*, namely ABHD14A and ABHD14B. Until our recent functional annotation of ABHD14B as a novel KDAC*^22, 23^*, both enzymes lacked any known function, despite their discoveries over two decades ago*^20, 21^*. Specifically, in this study, we provide a bioinformatics framework that helps unambiguously identify a sequence as ABHD14A or ABHD14B from various databases across diverse organisms. Further, we provide an updated list of ABHD14A and ABHD14B sequences from various organisms that possess them, along with a detailed phylogenetic analysis for these sequences for researchers interested in studying these enzymes. As a proof of concept, we also experimentally (biochemically) validate these findings by performing mutagenesis studies for a handful of residues for human ABHD14B, and find that there are residues distal to the active site (e.g. S75 for human ABHD14B), that might be regulate catalysis by possible allosteric mechanisms, a fact previously unknown for these enzymes. The knowledge and resources generated from this study will also be of biomedical importance, as it is now evident from several recent population-wide genetic association studies that dysregulation in ABHD14B expression has strong associations with different types of cancers*^37–42^*, while anomaly in ABHD14A expression is found to be associated with neurodegenerative conditions*^43–47^*, and even some cancers*^48–51^*.

## DATA AVAILABILITY STATEMENT

All the data that supports the findings of this study are available in the paper, and its associated supporting information, or are available from the corresponding authors upon reasonable request.

## CONFLICT OF INTEREST STATEMENT

The authors declare no conflicts of interest with the contents of this article.

## AUTHOR CONTRIBUTIONS

S.S.K. and M.S.M. conceived and supervised the project; K.V. performed all the biochemical studies on ABHD14B with assistance from M.Y.; G.R. performed the bioinformatics studies with assistance from K.V., S.G., and A.D.; S.S.K. acquired funding for this project. K.V., G.R., M.S.M. and S.S.K. wrote the paper with assistance from all authors.

## ACKNOWLEDGEMENTS

This work was supported by an EMBO Young Investigator Award and intramural funding from IISER Pune (to S.S.K.), and a Department of Biotechnology, Government of India grant to IISER Pune under the BICB Centre Scheme (BT/PR40262/BTIS/137/38/2022). K.V. and S.G. were supported by a Graduate Student Fellowship from IISER Pune and the Prime Minister’s Research Fellowship respectively. A.D. was supported by a DST-INSPIRE Scholarship.

## Notes

### Competing Interest Statement

The authors have declared no competing interest.

